# NanoLAS: a Comprehensive Nanobody Database with Data Integration, Consolidation, and Application

**DOI:** 10.1101/2023.09.27.559691

**Authors:** Shuchang Xiong, Zhengwen Liu, Xin Yi, Kai Liu, Bingding Huang, Xin Wang

## Abstract

Nanobodies, a unique subclass of antibodies first discovered in camelid animals, are composed solely of a single heavy chain’s variable region (VHH). Their significantly reduced molecular weight, in comparison to conventional antibodies, confers numerous advantages in the treatment of various diseases. As research and applications involving nanobodies expand, the quantity of identified nanobodies is also rapidly growing. However, the existing antibody databases are deficient in type and coverage, failing to satisfy the comprehensive needs of researchers and thus, impeding progress in nanobody research.

In response to this, we have amalgamated data from multiple sources to successfully assemble a new and comprehensive nanobody database. This database has currently included the latest nanobody data, provides researchers with an excellent search and data display interface, thus facilitating the progression of nanobody research and their application in disease treatment.

In summary, the newly constructed Nanobody Library and Archive System (NanoLAS) may significantly enhance the retrieval efficiency and application potential of nanobodies. We envision that NanoLAS will serve as an accessible, robust, and efficient tool for nanobody research and development, propelling advancements in the field of biomedicine.

**Database URL:** https://www.nanolas.cloud

## I. Introduction

Nanobodies, a unique subclass of antibodies discovered in camelid animals [1], are composed solely of a variable region (VHH), contributing to their compact structure and significant therapeutic advantages. The diverse properties of nanobodies underpin their broad application potential in biological research and disease treatment [2,3]. Nanobodies exhibit high specificity, solubility, stability, and antigen affinity with low toxicity and immunogenicity [4]. Furthermore, the small size allows nanobodies superior tissue penetration [5,6], making them advantageous in disease treatment and molecular imaging [7-9]. Nanobodies’ properties and smaller size enable effective tissue penetration, easy engineering, multimeric structures generation, and application in diverse fields, including cancer treatment and COVID-19 drug development [10-17].

The continuous advancement in nanobody research and applications has led to a rapid accumulation of nanobody data over recent years. Current databases, such as PDB, INDI, IMGT, Sdab-db, among others, house vast volumes of nanobody information. However, these databases may fall short in terms of data type and coverage.

Furthermore, the heterogeneity, inconsistency, and lack of interoperability of data across different databases pose additional challenges for researchers. Each database follows its unique data format and structure, obliging researchers to invest substantial time and effort in data processing and integration when using multiple databases. Some databases do not even offer a user-friendly interface, complicating and prolonging the data query and analysis process.

To address these limitations, we proposes the creation of a new nanobody database—NanoLAS. This initiative aims to satisfy the scientific community’s need for a more comprehensive and in-depth understanding of nanobodies. NanoLAS will integrate and standardize nanobody data from diverse databases, offer a user-friendly, efficient, and interactive query and analysis platform, and facilitate the further development of nanobody research.

## II. METHODS

### A. Data Collection

In the construction of our nanobody database, we have sourced data from multiple publicly accessible bioinformatics databases in the *Figure 1*. Given that each database employs unique information formats and content, it’s necessary to carefully process and convert this information specifically to ensure uniformity.

**Figure 1.**
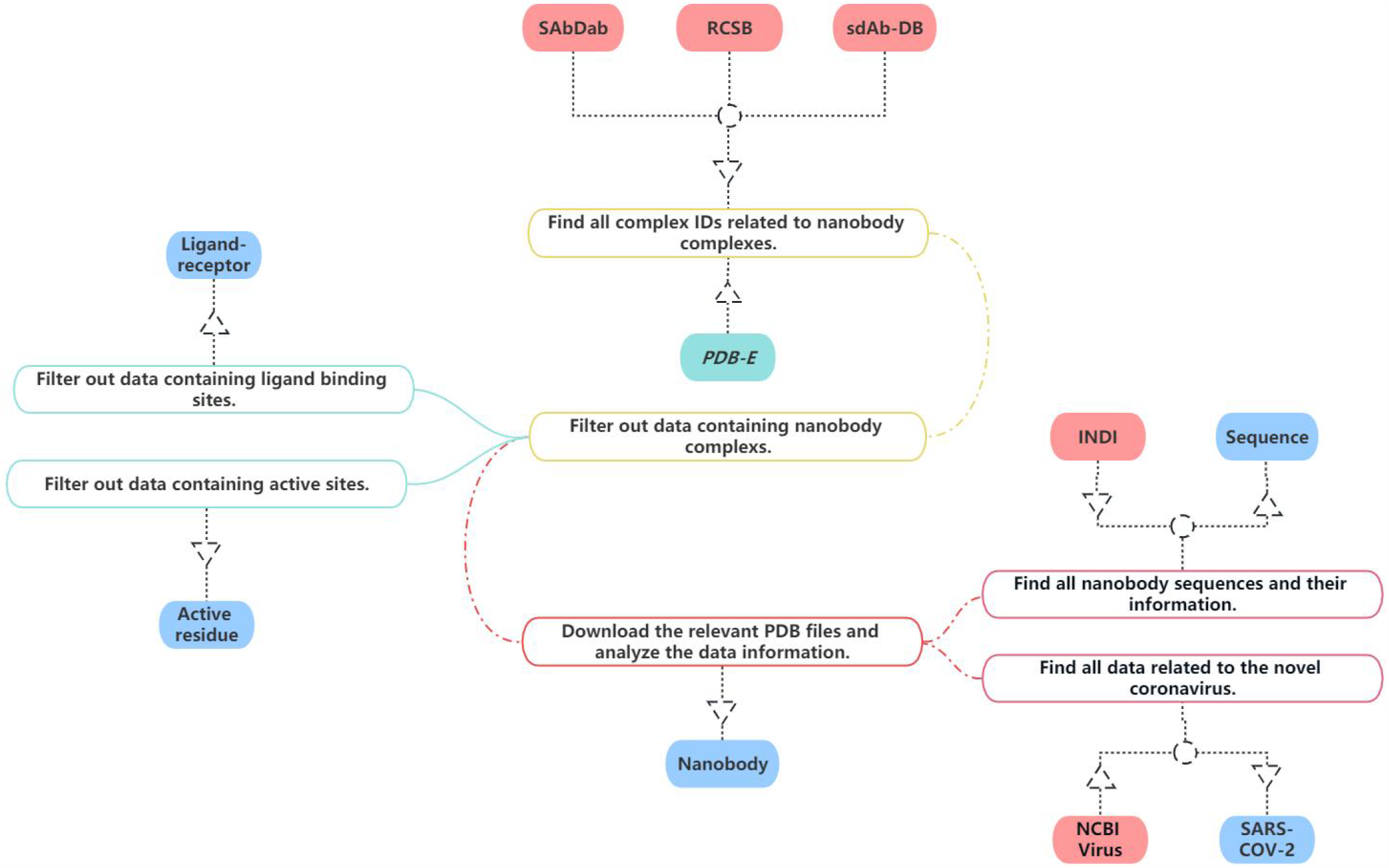
The process of data collection and processing of NanoLAS database.

For our project, we have gleaned all relevant nanobody protein structure information from the RCSB PDB [18]. Our selection and extraction process were based on a comprehensive set of screening criteria such as protein function, source, resolution, and publication date. This approach ensured that our collected data is representative and aligns with our research needs. Consequently, we successfully amassed a substantial amount of nanobody structure information, which forms a solid foundation for our database.

Sdab-db [19] provides detailed and meticulously verified single-domain antibody sequences, sources, structures, and associated biological information.We have selected and extracted nanobody-related data from Sdab-db, comprising sequences, structures, affinities, sources, and related biological information of single-domain antibodies.

Opig-SABDAB [20] gathers detailed records of the source, sequence, three-dimensional structure, and other biological information of nanobodies. We extracted relevant nanobody data from Opig-SABDAB. These data include but are not limited to the amino acid sequence, structural model, source information, and corresponding biological functions of nanobodies. Through the integration and utilization of these data, we have established a comprehensive and high-quality nanobody dataset in NanoLAS to support the research and application of nanobodies.

We extracted relevant SARS-CoV-2 data from the NCBI Virus database [21]. Through the integration of this data, we enriched the SARS-CoV-2 entry in the NanoLAS database, providing a robust resource to support research and applications focusing on SARS-CoV-2.

PDB-E [22] provides a variety of tools and services that allow users to search for and analyze structures available in the PDB archive. Some of its features include advanced search capabilities, analyses of macromolecular structures, sequence and structure alignment tools, and links to other resources for further analysis. We utilize these tools and enriched the Ligand-receptor and active residue entries in NanoLAS database.

Through the bulk download feature in INDI [23], we analyzed all nanobody sequences and integrated this data with the sequences gathered from RCSB, SAbDab, and sdAb-DB. As a result, we have constructed a comprehensive nanobody sequence entry within our NanoLAS database.

### B. Website Construction

In the construction of NanoLAS website (*Figure 2*), we utilized a modern technology stack for optimal efficiency and stability. The Vue.js framework, HTML, CSS, and JavaScript were employed for front-end development, creating user-friendly interfaces. Java Spring Boot was utilized for back-end functions, while MySQL was selected for its powerful data processing capabilities. For molecular structure visualization, we used 3Dmol.js [24], a JavaScript library. The system operates based on the HTTP protocol, where user-initiated front-end requests are processed by the backend and the corresponding results are rendered on the frontend.

**Figure 2.**
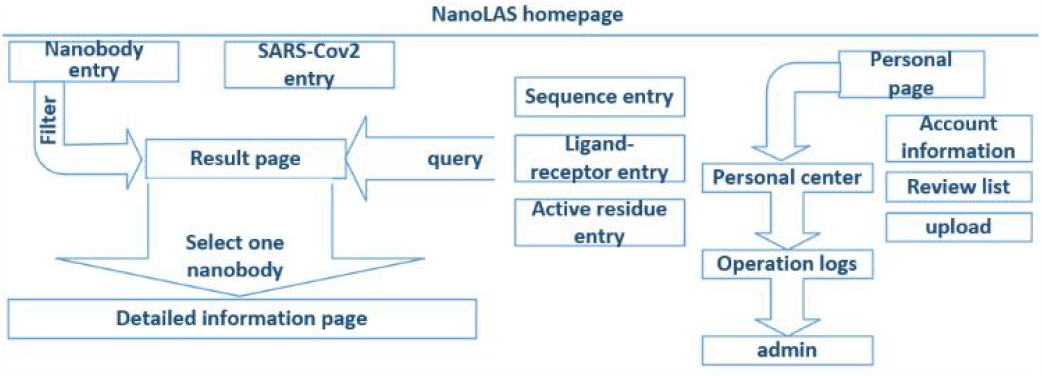
The architecture of NanoLAS website. iCAN website includes five search entries (left part) and a interface for users (right part).

## III. Results

### A. Data Overview

In the process of data integration, we have tried to include detailed information for each antibody sequence, such as source species, CDR region lengths, etc.

For organism sources, the database contains nanobody data from wide variety of species, and lama glama occupies the majority.

For sequences, we compared the length of CDR1∼3 in NanoLAS database. While CDR1 and CDR2 do not show obvious length variations, CDR3 performs the most variable portion. CDR3 lengths span a wide range from 5 to 28 amino acids, with lengths 14, 16, 17, and 21 being the most represented in the database (*Figure 3*).

**Figure 3.**
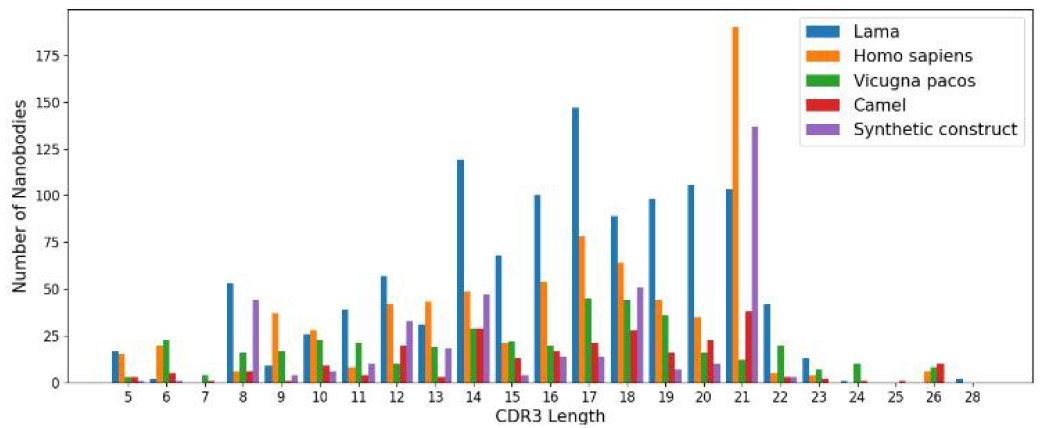
The distribution of CDR3 lengths in the NanoLAS database by source organism.

### B. Website Features and Operation Interface

The NanoLAS database can be accessed at http://www.nanolas.cloud, and we also maintain a project repository on GitHub, facilitating developers and researchers for deep participation and contribution.

#### 1) Search, Filter, and Display of Nanobodies (Figure 4)

NanoLAS allows users to flexibly retrieve nanobody data in the database by providing various filters and search tools. Users can operate through four main entrances or the search box at the top. NanoLAS provides the following multiple retrieval methods for various nanobody sequences in the database:

**Figure 4.**
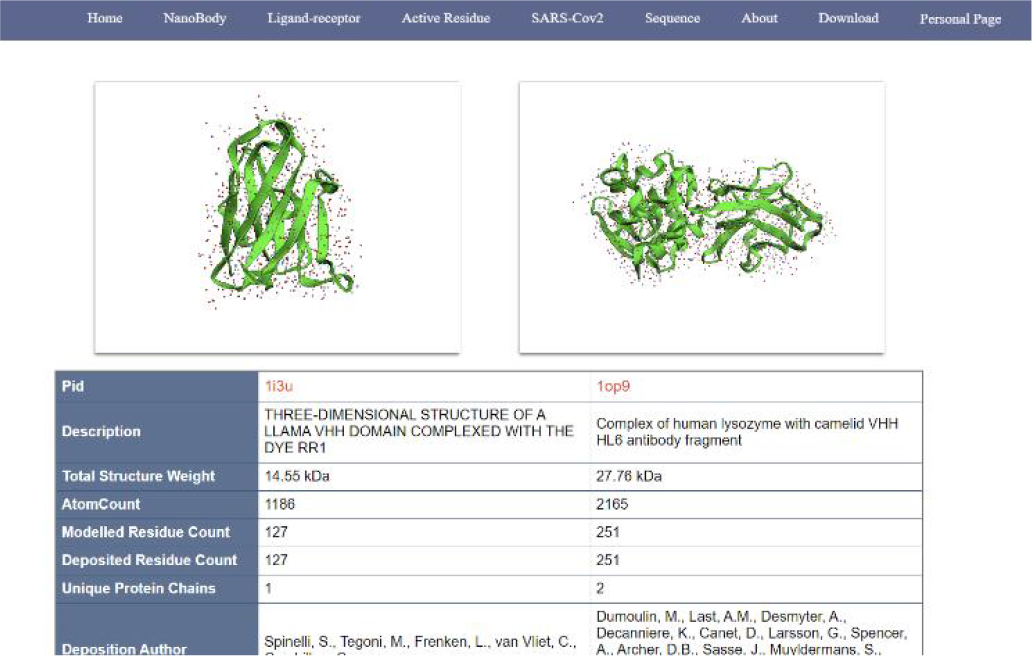
An example of information page of nanobodies (comparable/docking view).

- Identifier of the nanobody source database, such as PDB ID.
- Search by nanobody amino acid sequence.
- Search by CDR region.
- Year, source organism, etc.
- Ligand receptor.
- Active residues.

After retrieving the target antibody, users can choose different styles to observe the 3D structure, possibly using 3dmol. The search results also include pid, total structure weight, total number of atoms, number of residues, description, publication date, etc. The search results also contain a download link for the pdb file, allowing users to download the pdb file for more detailed operations and analysis.

1. *Comparison and Analysis of Nanobodies:* NanoLAS provides users with the function of comparing and analyzing two different antibodies. If you need to compare two specific nanobodies, users can set the parameters in the search box. Through 3Dmol.js, we can intuitively compare the similarities and differences between different nanobodies in the 3D view and understand the structure of various nanobodies.
2. *User Data Upload:* We realize that there may be new nanobody data in the actual research and application process. Therefore, to comprehensively and meticulously collect nanobody data, NanoLAS specially provides a data submission page for nanobodies. On the nanobody data submission page, users can submit their nanobody information, such as PID, description, publication date, total number of atoms, etc. At the same time, to ensure the accuracy and reliability of the submitted data, we have set up a data review process. After users submit their nanobody information, we will conduct a data review. Only after the review, these data will be added to the database. We welcome and encourage all scholars and researchers to actively share their nanobody data, which will help promote the progress of nanobody research.

## IV. Discussion

As a brand-new database specifically collecting nanobody information, NanoLAS has several advantages (*Figure 5)*. First, our database contains nanobody data from multiple sources, which is extensive and diverse, and can meet the search needs of different users. Although the structural data of nanobodies have been included in databases such as RCSB, IMGT/3Dstructure-DB, and others, NanoLAS’s handling and presentation of data is more humanized and easier for users to retrieve and understand. Secondly, NanoLAS provides a 3D view function, allowing users to intuitively view the spatial structure of nanobodies, which is very helpful for studying and understanding the structure and function of nanobodies. Furthermore, our comparative analysis function allows users to intuitively compare the differences between two or more nanobodies, assisting researchers in making scientific judgments. Finally, NanoLAS’s interface design is beautiful and easy to operate, providing users with a comfortable user experience.

**Figure 5.**
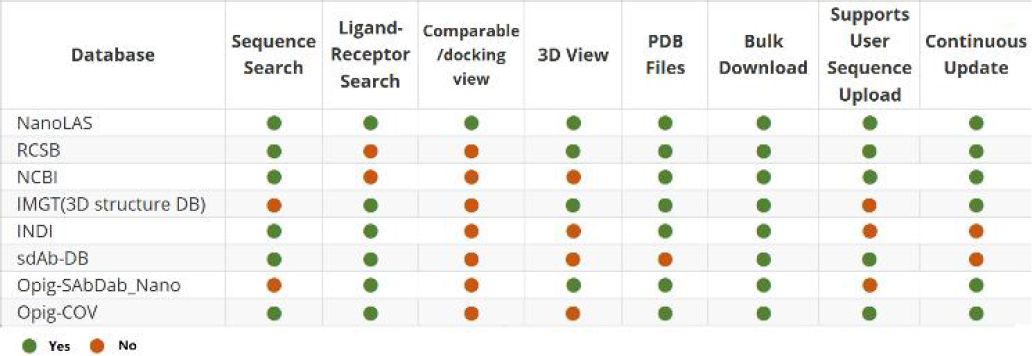
Comparison of NanoLAS and other databases.

With the progress of scientific research, nanobody sequence data is rapidly increasing and updating, it’s important to note that the success of the database will depend on its continuous update and maintenance, as well as the addition of new features and tools based on user feedback and needs. and the NanoLAS database needs to take measures to regularly increase the collection and update to maintain the timeliness of the collected data.

## V. Conclusion

In this project, we successfully developed the nanobody database NanoLAS. It integrates nanobody data from multiple sources, provides a user-friendly interface, allowing users to easily query, analyze, and visualize nanobody data, thereby improving research efficiency.

To continuously optimize and develop NanoLAS, we sincerely invite and look forward to various feedback and suggestions from users. We will strive to improve and expand NanoLAS to better serve the research and application field of nanobodies. For suggestions or queries, please reach out to us through our contact page.

## Notes

### Competing Interest Statement

The authors have declared no competing interest.

